# Held out wings RNA binding activity in the cytoplasm during early spermatogenesis

**DOI:** 10.1101/2022.05.19.492257

**Authors:** Michaela Agapiou, Tayah Hopes, Fruzsina Hobor, Amanda Bretman, Thomas A. Edwards, Julie L. Aspden

## Abstract

Held out wings (HOW) is an RNA-binding protein essential for spermatogenesis in *Drosophila melanogaster*. HOW is a signal transduction and activation of RNA (STAR) protein, regulating post-transcriptional gene expression. The characteristics of RNA-binding by the conserved short cytoplasmic isoform, HOW(S), are unknown. *In vivo* RIP-seq identified 121 novel transcripts bound by HOW(S) in germ stem cells and spermatogonia, many with signal transduction functions. (A/G/U)CUAAC motifs were enriched in 3’-UTRs and GCG(A/U)G in 5’-UTRs. HOW binds with high-affinity to sites containing CUAAC motifs from *lola* and *hipk* mRNAs. This study provides new insight into STAR protein-RNA interactions and functions in spermatogenesis.

## Introduction

Post-transcriptional gene regulation plays an essential role in stem cells contributing to pluripotency, differentiation and self-renewal. Molecular signals contribute to germ stem cell regulation through a variety of RNA processes controlled by a network of RNA-binding proteins including the signal transduction and activation of RNA (STAR) protein family. STAR proteins characteristically contain a single RNA-binding STAR domain, composed of one maxi-K-homology (KH) domain, and one or two flanking regions (Vernet & Artzt, 1997). As a result of their functions in signal transduction pathways, the family contributes to the regulation of gene expression during key stages of development. For example, the STAR protein Quaking (QKI) regulates RNA processing during gametogenesis and myelination in mammals (Kondo et al., 1999). RNA-binding proteins in general are important throughout spermatogenesis, including for stem cell maintenance and meiosis, across a variety of organisms (Legrand & Hobbs, 2018).

Held out wings (HOW) is the ortholog of QKI in *Drosophila melanogaster*. HOW is essential but partial loss of function mutants exhibit a variety of phenotypes including defective wing development, glial maturation, tendon maturation and sterility (Baehrecke, 1997; Edenfeld et al., 2006; Monk et al., 2010; Nabel-Rosen et al., 1999). HOW regulates RNA processing, including pre-mRNA splicing, localisation and mRNA degradation (Graindorge et al., 2013; Nabel-Rosen et al., 2002; Rodrigues et al., 2012). There are multiple protein isoforms of HOW, produced through alternative splicing, all of which contain the STAR domain but differing at the C-terminus (Larkin et al., 2021). The different isoforms of QKI and HOW are differentially localised within cells, and the shortest isoforms, QKI-6 and HOW(S), are predominantly cytoplasmic, where their function is unknown. The HOW and QKI ortholog in *C. elegans*, GLD-1, has a single cytoplasmic isoform that regulates mRNA translation in germ cells (Jan et al., 1999). Mirroring QKI’s expression in mammals, HOW is expressed in *D. melanogaster* testis, and it is important during spermatogenesis (Monk et al., 2010). In germ cells, HOW’s expression is unusual and highly restricted to the earliest stages of gametogenesis, where it contributes to the maintenance of germ stem cells (GSCs); without HOW, GSCs in the testis do not survive (Monk et al., 2011; Monk et al., 2010). Expression of HA-tagged HOW(S) in the testis indicates that this isoform is restricted to the cytoplasm in germ cells (Monk et al., 2010). The restricted expression of HOW during spermatogenesis suggests a specific function in regulating the balance between stem cell maintenance versus the proliferation and differentiation of spermatogonia.

The majority of work on HOW thus far has focused on HOW(L) and its role in nuclear RNA processing events. Current understanding of HOW RNA binding comes from a handful of mRNAs (e.g. *dpp, miple1, bam*), which contain 3’-UTR ACUAA motifs typically bound by HOW(L) (Israeli et al., 2007). The only mRNA previously known to be bound specifically by HOW(S) rather than HOW(L) is *dgrasp* in oocytes (Giuliani et al., 2014). An optimal binding preference has yet to be characterised for HOW but a consensus of NCUAACN has been generated from *in vitro* binding experiments (Ray et al., 2013). Here, we sought to globally identify novel mRNA targets of HOW in the testis, specifically by HOW(S) in the cytoplasm, to understand the binding characteristic of this interaction *in vivo*.

## Results & Discussion

### Expression and pull-down of cytoplasmic HA-tagged HOW(S) in testis germ cells

To understand the role of HOW(S) and the RNAs it binds in the cytoplasm, HA-tagged HOW(S) was expressed in the early stages of spermatogenesis in *Drosophila melanogaster* testes. This was achieved by using the UAS/GAL4 system, whereby HOW(S)-HA expression was driven by *nanos*-GAL4, and thus, HOW(S)-HA was expressed in GSCs and early spermatogonia (Fig. 1A). This HA-tagged HOW(S) is localised to the cytoplasm of these cells (Fig. 1A), recapitulating the presence of cytoplasmic HOW in these specific germ cells (Monk et al., 2011; Monk et al., 2010). We were able to successfully pull down HOW(S)-HA from the lysates of these testes using anti-HA beads (Fig. 1B). To isolate RNA bound to HOW(S), RNP-IPs were performed from large scale testes lysates in triplicate (Fig. 1C and Supplemental Fig. 1A-C). RNA from the HOW(S)-HA pull-down was purified along with RNA from input testis lysates and RNA from pull-downs performed from the *nanos*-GAL4 parent testes, which had no HA-tagged HOW(S) (Supplemental Fig. 1D). These RNA samples then underwent RNA-seq.

**Figure 1:**
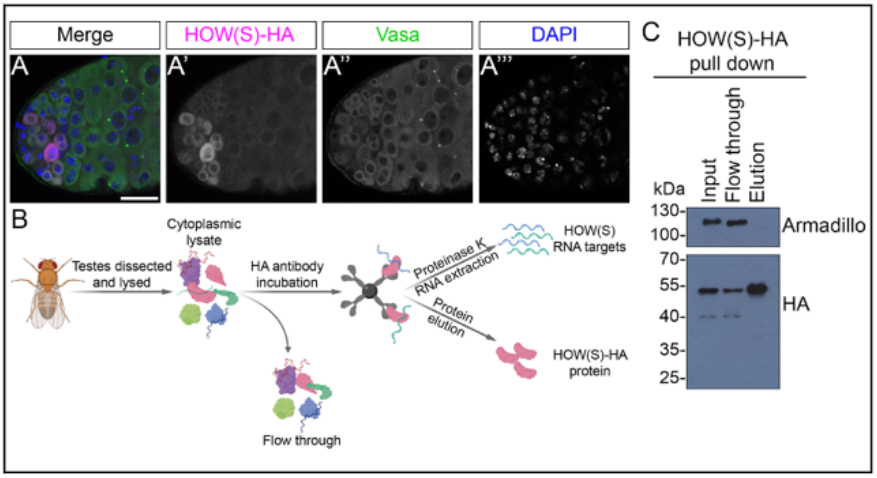
Expression and pull-down of HA-tagged HOW(S) from cytoplasm of germ cells. A) Confocal images showing cytoplasmic expression of HOW(S)-HA (magenta)in GSCs and spermatogonia, driven by the nanos-GAL4 line. Germ cells are positive for vasa (green), and blue is DAPI. B) Schematic of HOW(S)-HA RIP-seq. C) Western blot showing specific depletion of HOW(S)-HA from testis the input lysate and enrichment in the elution using anti-HA beads.

### HOW(S) bound RNAs are enriched for signalling functions

To identify which RNAs were specifically bound to HOW(S)-HA, differential transcript enrichment analysis was performed on RNAs in HOW(S)-HA pulldown compared to those present in input lysate (Fig. 2A, Supplemental Table 1, Supplemental Fig. 2A-B). The majority of transcripts showed no enrichment, having a log^2^(Fold Change) between -0.5 and 0.5 (Fig. 2B). We defined enriched transcripts as those with a log^2^(Fold Change) ≥ 1 and an FDR-corrected *p*-value ≤ 0.05. Through this analysis we identified 403 transcripts enriched in the HOW(S)-HA pull-down RNA compared with input RNA. Non-specific background RNAs enriched in the parental *nanos*-GAL4 RIP-seq sample (with no HOW(S)-HA present) were then subtracted, leaving those RNA transcripts specifically bound to HOW(S)-HA. 121 mRNAs were identified as specifically enriched by HOW(S) RIP-seq (Fig. 2C – green dots). To understand the role of the RNAs bound by HOW(S), GO analysis was performed on the genes corresponding to these HOW(S) bound mRNAs. This revealed an enrichment for genes with functions in cell signalling, specifically in cell communication and signal transduction e.g. *hipk* (homeodomain interacting protein kinase) and *CycG* (cyclin G) (Fig. 2D, Supplemental Fig. 2C). This is entirely consistent with HOW’s membership of the STAR family of proteins.

**Figure 2:**
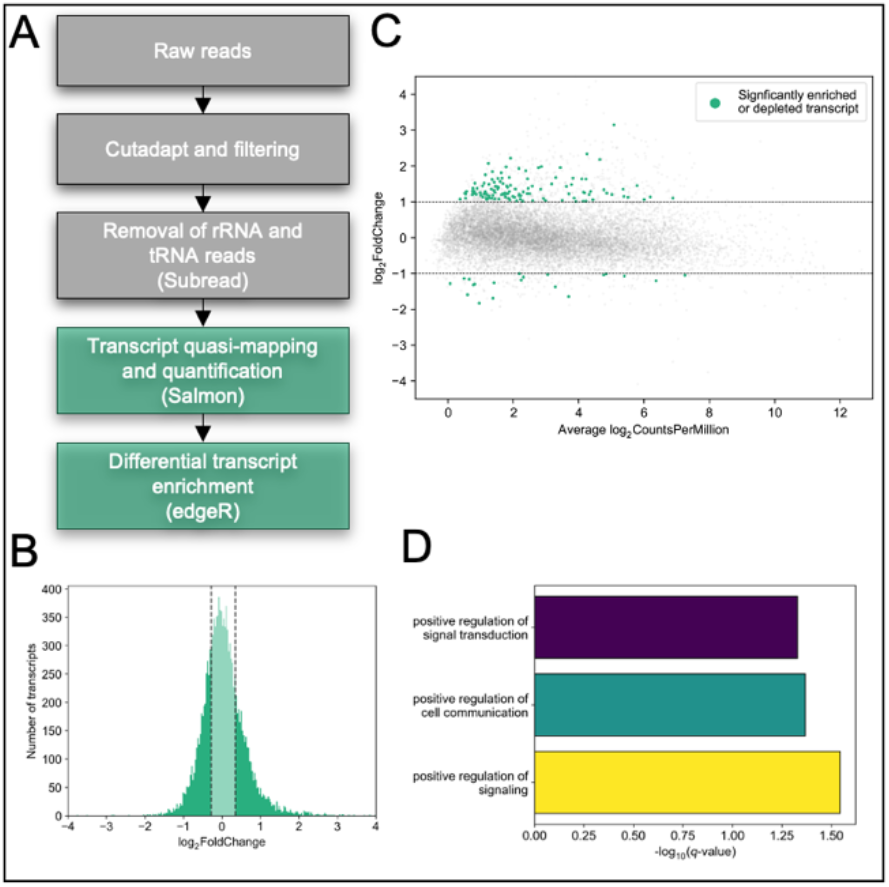
Enrichment of HOW(S) bound RNA, with role in signal transduction. A) Pipeline to identify RNAs specifically bound to HOW(S). B) Histogram from differential enrichment analysis of pull-down versus input, middle quartiles are shaded in lighter green. C) Scatter plot from differential transcript analysis. Significantly enriched or depleted transcripts are those with log^2^(Fold Change) of ≥ 1 or ≤ -1 (dotted lines), and an FDR-corrected p-value ≤ 0.05, additionally non-specific background RNAs enriched in the parental nanos-GAL4 RIP-seq sample were subtracted. Specifically enriched or depleted RNAs from the HOW(S) RIP-seq are highlighted in green. D) GO term analysis reveals over-representation of signaling related terms in transcripts bound by HOW(S).

HOW’s ortholog QKI is a known regulator of circRNA processing (Conn et al., 2015), and while our method was not optimised for circRNA detection, further analysis revealed a small number of circRNAs in our dataset, some of which HOW(S) may bind (Supplemental Fig. 2D and Supplemental Table 2).

### CUAAC motif enriched in 3’-UTRs of HOW(S) bound RNAs

To determine the nature of HOW(S) binding to the 121 transcripts identified by RIP-seq, motif analysis was performed on these mRNA targets. Within the 3’-UTRs the most enriched motif was found to be (A/G/U)CUAAC (Fig. 3A), with 46% of the HOW(S)-HA bound mRNAs containing this motif (Fig. 3B). (A/G/U)CUAAC is very similar to the other previously identified HOW binding sequence in *stripe* and *dpp* mRNAs: ACUAA (Israeli et al., 2007). Rather than characterising a small number of HOW targets and identifying individual binding sites, the (A/G/U)CUAAC motif generated here is from a much larger number of RNAs bound by HOW(S) in the cytoplasm.

**Figure 3:**
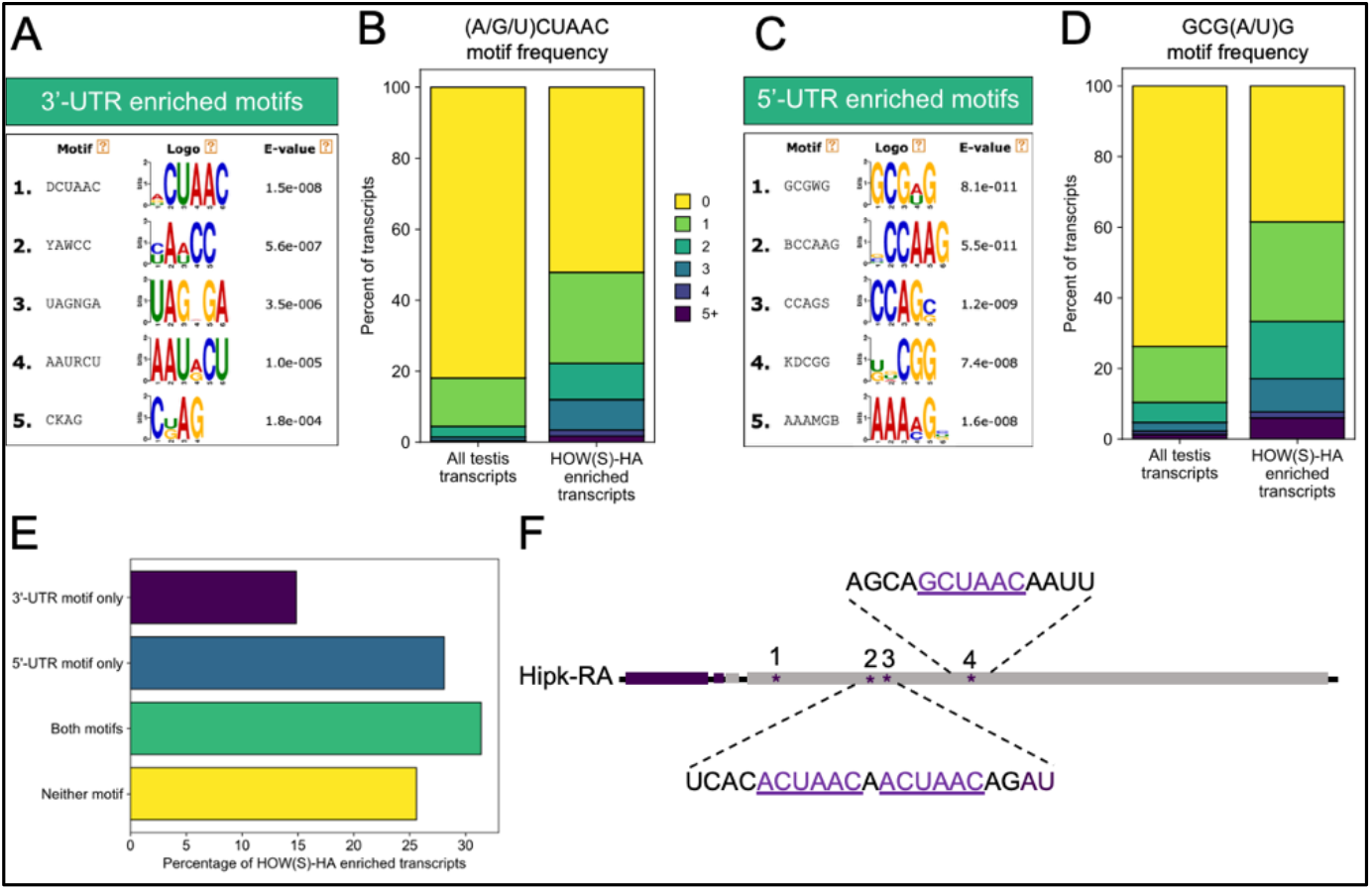
Enrichment of CUAAC motifs in HOW(S) bound RNAs. A) Motifs found to be enriched in 3’-UTRs of mRNAs bound by HOW(S). B) Frequency at which (A/G/U)CUAAC motif identified in 3’-UTR of HOW(S) bound transcripts compared to the 3’-UTRs of all testis expressed transcripts. C) Motifs found to be enriched in 5’-UTRs of mRNAs bound by HOW(S). D) Frequency at which GCG(A/U)G motif identified in 5’-UTR of HOW(S) bound transcripts compared to the 5’-UTRs of all testis expressed transcripts. E) Percentage of transcripts bound by HOW(S) that had either of the top UTR motifs, both, or neither in their respective UTRs. F) Schematic of the 3’ end of hipk transcript, purple is CDS, grey is UTR, and asterisks mark (A/G/U)CUAAC sites in the 3’-UTR.

Other distinct motifs were found enriched in the 5’-UTRs of HOW(S) bound mRNAs, including GCG(A/U)G (Fig. 3C), but these do not show similarity with previously identified HOW binding sites (Graindorge et al., 2013; Israeli et al., 2007; Ray et al., 2013). Although these are unlikely to be HOW(S) binding sites, because their sequence differs dramatically from the previously characterised ACUAA motif, these motifs may be important for HOW(S) RNA-protein complexes as they are highly enriched in the HOW(S) bound mRNAs – the GCG(A/U)G motif is present in 60% of HOW(S) bound mRNAs (Fig. 3D). Instead, these motifs may represent sequences that are bound to another RNA-binding protein (RBP) associated with HOW(S), since RIP-seq will identify both indirect and direct interactions. This type of co-binding has been demonstrated for HOW(L), which can interact with the RBP SXL, with the two proteins binding distinct sequences in the 5’-UTR of *msl-2* (Graindorge et al., 2013). We also observed that 31% of HOW(S) bound mRNAs contain enriched motifs both in their 5’- and 3’-UTRs (Fig. 3E), which indicates that HOW(S) molecules could bind in both UTRs of an mRNA transcript simultaneously, potentially regulating the mRNA in numerous ways. This has been shown for other RBPs during development e.g. SXL during sex determination (Duncan et al., 2006).

Further analysis of the HOW(S) bound transcripts that have the (A/G/U)CUAAC motif reveals that 21% of them contain multiple repeats of this motif, which is substantially higher than that found in all testis expressed transcripts (Fig. 3B). RBPs often bind RNAs in a modular manner, and HOW, along with many of the STAR proteins, dimerise (Nir et al., 2012). Thus, multiple binding sites in HOW(S) bound transcripts could indicate a modular binding mode. This pattern is also seen for the top 5’-UTR enriched motif GCG(A/U)G (Fig. 3D), indicating that this too may represent a modular RBP binding region.

One such mRNA which has multiple predicted binding sites is *hipk*. Hipk protein functions in multiple signalling pathways, including Notch (Lee et al., 2009), Hippo (Chen & Verheyen, 2012; Poon et al., 2012), and the JAK-STAT pathway (Tettweiler et al., 2019) and is essential for germline development in *C. elegans* and mouse spermatogenesis (Berber et al., 2013; Crapster et al., 2020). *Hipk-RA* transcript (FBtr0072552) was identified as enriched in the HOW(S)-HA RIP-seq and contains four (A/G)CUAAC sites within its 3’-UTR (Fig. 3F), therefore likely represents a key HOW(S) target with downstream roles in signalling pathways essential in maintaining the stem cell niche and cell fate.

### Recombinant HOW-STAR domain binds enriched motifs within HOW(S) bound RNAs with high affinity

To test the binding of HOW to the motifs identified as enriched from RIP-seq we recombinantly expressed HOW’s STAR domain (Fig. 4A, Supplemental Fig. 3A-C) and performed fluorescence anisotropy (FA) binding assays. Three different mRNAs that we identified as being bound by HOW(S) in GSCs and early spermatogonia were selected to test HOW binding to sequences from within these transcripts that were identified as highly similar to the enriched motifs.

**Figure 4:**
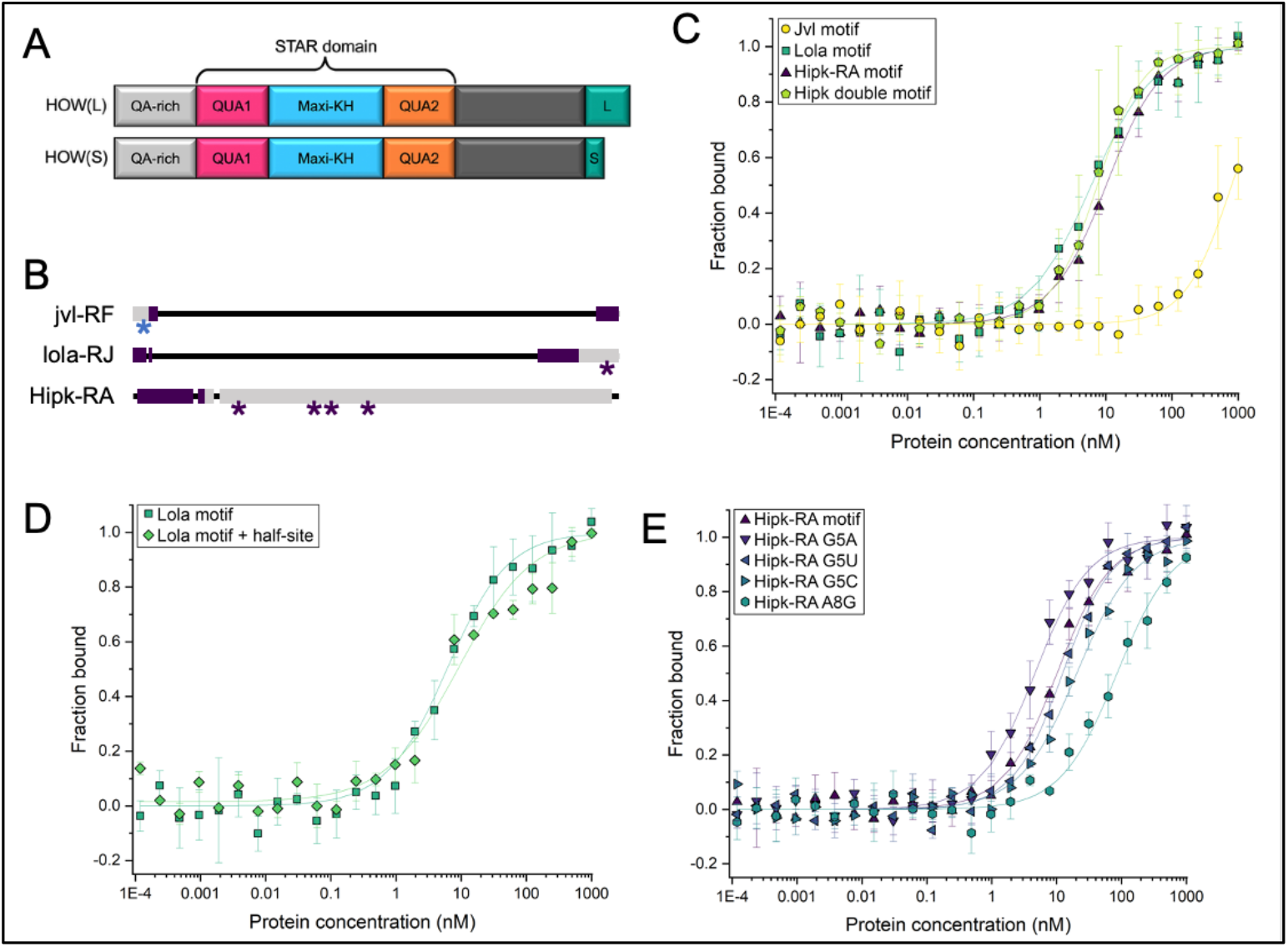
HOW KH domain binds to HOW(S) targets with high affinity and specificity. A) Schematic of two HOW protein isoforms, indicating recombinant protein STAR domain expressed. B) Schematic of three transcripts bound by HOW(S): the 5’ end of jvl-RF, and 3’ ends of lola-RF and Hipk-RA. Asterisks mark the 5’-UTR (blue) and 3’-UTR (purple) motifs. Purple bars are CDS, grey is UTRs. C) FA binding plots from jvl motif, lola motif and Hipk-RA motif, as well as the Hipk double site. D) lola motif and lola motif + half-site. E) Mutations in the Hipk-RA motif.

A site was selected from the 5’-UTR of *jvl* transcript (FBtr0305694) that contained the most highly enriched 5’-UTR motif within HOW(S) bound mRNAs: GCGUG (Fig. 4B). FA analysis of this *jvl* RNA oligo revealed that HOW-STAR did not bind to this site in the low nanomolar range (Fig. 4C & Table 1). This suggests that although the GCG(A/U)G motif is highly enriched in the 5’-UTRs of HOW(S) bound mRNAs, it is unlikely to be an *in vivo* HOW(S) binding site. An alternative explanation could be that HOW(S) interacts with a second RBP, which provides the RNA-binding specificity for these interactions. One potential candidate is SLIRP1 protein, which has a binding motif that is highly similar to the one found in the 5’-UTR of HOW(S) bound RNAs: GCG(U/C)(G>A/C/U) (Ray et al., 2013) and is highly expressed in the testis.

**Table 1:**
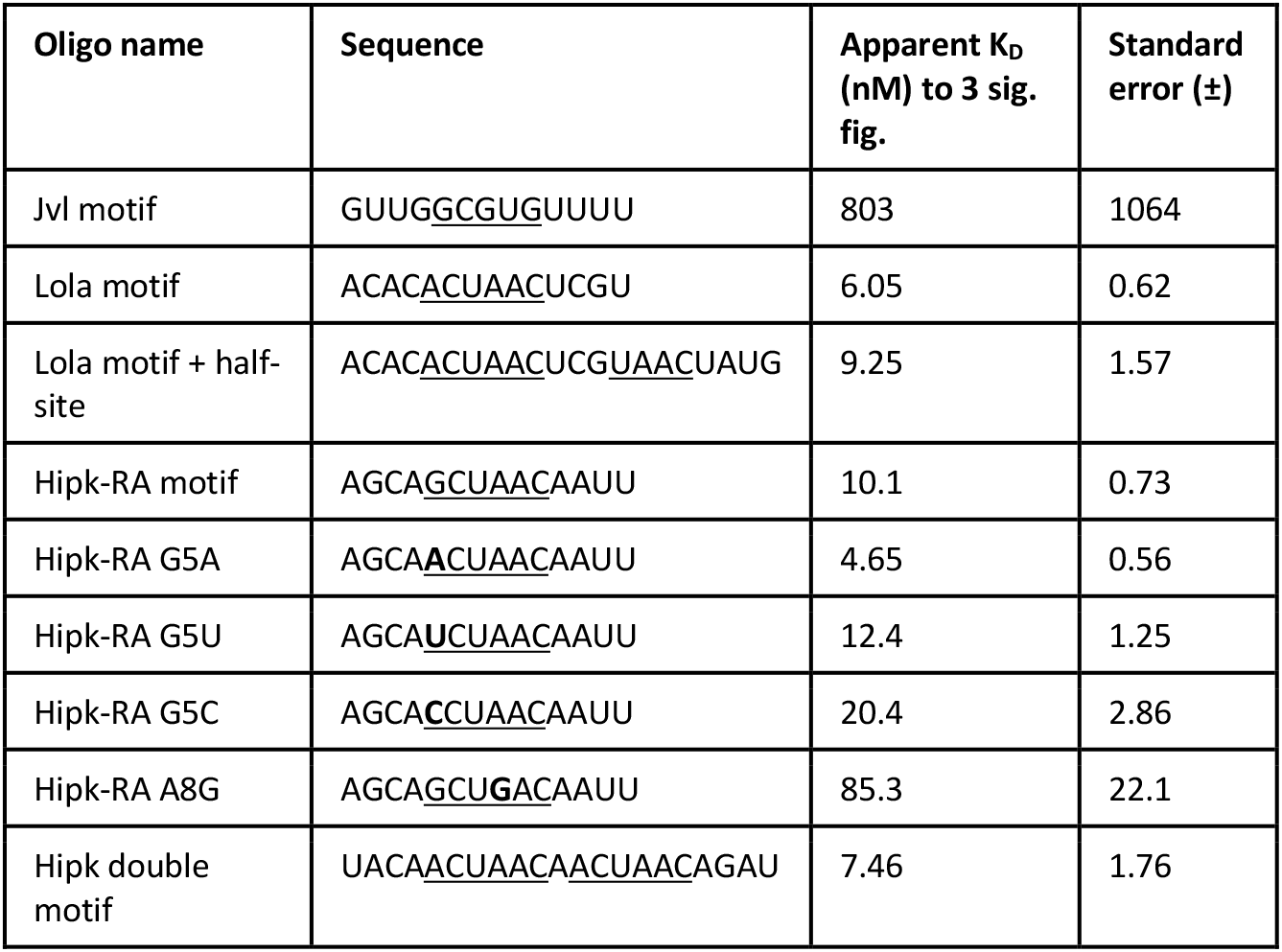
Summary of HOW binding affinities for HOW(S) predicted binding sites. Sequence of oligos based on the predicted binding sites within HOW(S) targets, along with apparent K_D_ and standard error calculated from FA. Sequences that match the top motifs from the motif enrichment analysis are underlined, and nucleotides mutated are in bold.

We also tested RNA oligos based on regions from *lola* and *hipk*, two transcripts bound by HOW(S), which contain sequences matching the 3’-UTR HOW(S) motif. The *lola* element contains an ACUAAC motif and HOW-STAR exhibits high affinity binding to this RNA oligo by FA (Fig. 4C, Supplemental Fig. 3D). Just downstream of this motif is a potential half-site and such half-sites have been suggested to enable stronger binding by STAR proteins (Galarneau & Richard, 2009). However, the RNA oligo including this additional half-site did not increase the affinity of HOW-STAR for the HOW binding site (Fig. 4D & Table 1). Therefore, we focused simply on HOW(S) binding sites with a complete motif.

Multiple sites were found in Hipk-RA mRNA that match the 3’-UTR enriched motif, so we characterised the binding of HOW(S) to RNA oligos containing the sequences of two of these regions (Fig. 3F). The first site contains GCUAAC and exhibits high affinity binding (10.1 +/- 0.73 nM) with HOW-STAR (Fig. 4C). Our motif enrichment analysis indicated that the first position of the top 3’-UTR motif, (A/G/C)CUAAC, appeared to be the most flexible (Fig 3A). Therefore, the G from this GCUAAC site was mutated to A, C or U, and binding tested to understand the effect on HOW-STAR binding affinity in the context of the same 14mer oligo used in these FA experiments. This revealed that there is a preference for adenosine at this site over guanosine, uridine and cytidine (Fig. 4E & Table 1). This is consistent with the motif enrichment analysis, where adenosine is more commonly found compared with other nucleotides (Fig. 3A). To show specificity of HOW binding to these NCUAAC motifs, the adenosine in the fourth position was mutated to a guanosine, which resulted in a large decrease (approximately 20 fold vs ACUAAC) in HOW-STAR binding affinity (Fig. 4E; Table 1). This is also consistent with the mutational analysis performed on HOW’s binding site in *stripe*’s 3’-UTR, which can be bound by either HOW(S) or HOW(L) (Israeli et al., 2007).

The second Hipk-RA site contains two HOW(S) motifs in close proximity in the 3’-UTR. The binding of HOW-STAR with this region, the Hipk ‘double site’, was of very similar affinity to the site only containing one CUAAC motif, suggesting that only a monomer of HOW is binding to this region, rather than dimer (Fig. 4C, Table 1). Other STAR proteins can bind as dimers and this may be the case for HOW but at more distant sites than those within this ‘double site’ (Table 1), although avidity effects may also be in play here.

Overall, we have identified 121 novel targets of cytoplasmic HOW(S) in testis germ cells, several of which are involved in signal transduction. Within these HOW(S) bound RNAs the (A/G/U)CUAAC motif is enriched in 3’-UTRs and GCG(A/U)G in 5’-UTRs. This 3’-UTR motif is similar to potential HOW binding sites identified in *stripe* and *dpp* mRNAs, as well the consensus sequences of other STAR proteins. The STAR domain of HOW has strong (nanomolar K^D^) affinity for RNA elements containing the CUAAC motif from *lola* and *hipk* mRNAs but not with GCG(A/U)G 5’-UTR motif. Together these results provide new insight into STAR protein-RNA interactions and potential importance to spermatogenesis. Given the importance of HOW to germ stem cells, understanding these novel cytoplasmic protein-RNA networks and their functional consequences will shed new light on how the balance of self-renewal and differentiation is regulated in stem cells.

## Methods

### Fly husbandry and stocks

Flies were kept in a 25°C humidified room with a 12:12 hour light:dark cycle and raised on 10ml standard sugar-yeast-agar medium (Bass et al., 2007). Crosses were generated by collecting unmated flies, then 5-10 flies of each sex were placed into a vial together with grains of active baker’s yeast. UAS-HOW-S-HA line generously gifted by Prof T. Volk, *nanos*-GAL4 line used is #64277 from BDSC.

### Immunofluorescence

Testes from 0-3 day old unmated flies were fixed with 4% paraformaldehyde and stained sequentially with anti-vasa then anti-HA. Slides were imaged using a Zeiss LSM880 Upright Confocal Microscope with the 40X oil-immersion objective and Zen imaging software. See the Supplemental Methods for details regarding tissue dissection, staining and antibodies.

### Ribonucleoprotein immunoprecipitation

1000 pairs of testes from 0–3 day old flies (mated or unmated) were dissected per sample. RIP pull-downs were adapted from (Keene et al., 2006), for full details see the Supplemental Methods.

### Sequencing and computational analysis

The RNA from the lysates and elutions of three HOW(S)-HA pull-downs and one *nanos*-GAL4 parental control pull-down was prepared using the Ribo-Zero rRNA Removal Kit (Illumina) followed by the TruSeq Stranded Total RNA Library Prep (Illumina), sequencing was 75 bp single-end. Differential transcript enrichment was carried out using edgeR, and motif enrichment with DREME. See Supplemental Methods for details regarding sequencing, filtering, mapping, and analyses.

### Data availability

RIP-seq data have been deposited to the Gene Expression Omnibus with accession ID: GSE201319.

### Protein expression and purification

Residues 72–266 of *D. melanogaster* HOW (the STAR domain) was expressed in BL21(DE3) cells with a His^6^-GST tag and purified via affinity chromatography and size exclusion chromatography. See Supplemental Methods for additional protein expression and purification details.

### Fluorescence anisotropy

RNA oligonucleotides were 3’ labelled with 6-carboxyfluorescein and mixed with purified STAR protein, the top protein concentration was 1 nM. See Supplemental Methods for assay and calculation details.

## Acknowledgments

We thank the Next Generation Sequencing facility, at St James University Hospital, Leeds, UK for performing Next Generation Sequencing. Parts of this work were undertaken on ARC3, part of the High Performance Computing facilities at the University of Leeds, UK. We thank the Protein Production Facility, especially Dr Brian Jackson. We also thank the Imaging Facility, Faculty of Biological Sciences, University of Leeds, UK for their assistance in confocal microscopy. We also thank the Biomolecular Mass Spectrometry Facility, University of Leeds, UK for their assistance.

## Author contributions

MA conceived, designed and performed experiments for the study. FH performed experiments for the study. TH performed experiments for the study. AB conceived the work, interpreted data, drafted and revised the manuscript. TE conceived the work, interpreted data, drafted and revised the manuscript. JLA conceived the work, interpreted data, drafted and revised the manuscript. All authors contributed to manuscript writing, revision and have approved the submitted version.

## References

Baehrecke, E. H. (1997). who encodes a KH RNA binding protein that functions in muscle development. Development, 124(7), 1323–1332.

Bass, T. M., Grandison, R. C., Wong, R., Martinez, P., Partridge, L., & Piper, M. D. W. (2007). Optimization of dietary restriction protocols in Drosophila. Journals of Gerontology - Series A Biological Sciences and Medical Sciences, 62(10), 1071–1081. https://doi.org/10.1093/gerona/62.10.1071

Berber, S., Llamosas, E., Thaivalappil, P., Boag, P. R., Crossley, M., & Nicholas, H. R. (2013). Homeodomain interacting protein kinase (HPK-1) is required in the soma for robust germline proliferation in <i>C. elegans</i>. Developmental Dynamics, 242(11), 1250–1261. https://doi.org/10.1002/dvdy.24023

Chen, J., & Verheyen, E. M. (2012). Homeodomain-interacting protein kinase regulates yorkie activity to promote tissue growth. Current Biology, 22(17), 1582–1586. https://doi.org/10.1016/j.cub.2012.06.074

Conn, S. J., Pillman, K. A., Toubia, J., Conn, V. M., Salmanidis, M., Phillips, C. A., … Goodall, G. J. (2015). The RNA binding protein quaking regulates formation of circRNAs. Cell, 160(6), 1125–1134. https://doi.org/10.1016/j.cell.2015.02.014

Crapster, J. A., Rack, P. G., Hellmann, Z. J., Le, A. D., Adams, C. M., Leib, R. D., … Chen, J. K. (2020). HIPK4 is essential for murine spermiogenesis. eLife, 9, e50209–e50209. https://doi.org/10.7554/eLife.50209

Duncan, K., Grskovic, M., Strein, C., Beckmann, K., Niggeweg, R., Abaza, I., … Hentze, M. W. (2006). Sex-lethal imparts a sex-specific function to UNR by recruiting it to the msl-2 mRNA 3’ UTR: translational repression for dosage compensation. Genes Dev, 20(3), 368–379. https://doi.org/10.1101/gad.371406

Edenfeld, G., Volohonsky, G., Krukkert, K., Naffin, E., Lammel, U., Grimm, A., … Klämbt, C. (2006). The splicing factor crooked neck associates with the RNA-binding protein HOW to control glial cell maturation in Drosophila. Neuron, 52(6), 969–980. https://doi.org/10.1016/j.neuron.2006.10.029

Galarneau, A., & Richard, S. (2009). The STAR RNA binding proteins GLD-1, QKI, SAM68 and SLM-2 bind bipartite RNA motifs. BMC Molecular Biology, 10(47), doi:10.1186/1471-2199-1110-1147. https://doi.org/10.1186/1471-2199-10-47

Giuliani, G., Giuliani, F., Volk, T., & Rabouille, C. (2014). The <i>Drosophila</i> RNA-binding protein HOW controls the stability of <i>dgrasp</i> mRNA in the follicular epithelium. Nucleic Acids Research, 42(3), 1970–1986. https://doi.org/10.1093/nar/gkt1118

Graindorge, A., Carré, C., & Gebauer, F. (2013). Sex-lethal promotes nuclear retention of <i>msl2</i> mRNA via interactions with the STAR protein HOW. Genes & Development, 27(12), 1421–1433. https://doi.org/10.1101/gad.214999.113

Israeli, D., Nir, R., & Volk, T. (2007). Dissection of the target specificity of the RNA-binding protein HOW reveals dpp mRNA as a novel HOW target. Development, 134(11), 2107–2114. https://doi.org/10.1242/dev.001594

Jan, E., Motzny, C. K., Graves, L. E., & Goodwin, E. B. (1999). The STAR protein, GLD-1, is a translational regulator of sexual identity in Caenorhabditis elegans. The EMBO Journal, 18(1), 258–269.

Keene, J. D., Komisarow, J. M., & Friedersdorf, M. B. (2006). RIP-Chip: The isolation and identification of mRNAs, microRNAs and protein components of ribonucleoprotein complexes from cell extracts. Nature Protocols, 1(1), 302–307. https://doi.org/10.1038/nprot.2006.47

Kondo, T., Furuta, T., Mitsunaga, K., Ebersole, T. A., Shichiri, M., Wu, J., … Abe, K. (1999). Genomic organization and expression analysis of the mouse qkI locus. Mammalian Genome, 10(7), 662–669. https://doi.org/10.1007/s003359901068

Larkin, A., Marygold, S. J., Antonazzo, G., Attrill, H., Dos Santos, G., Garapati, P. V., … Consortium, F. (2021). FlyBase: updates to the Drosophila melanogaster knowledge base. Nucleic Acids Res, 49(D1), D899–D907. https://doi.org/10.1093/nar/gkaa1026

Lee, W., Andrews, B. C., Faust, M., Walldorf, U., & Verheyen, E. M. (2009). Hipk is an essential protein that promotes Notch signal transduction in the Drosophila eye by inhibition of the global co-repressor Groucho. Developmental Biology, 325(1), 263–272. https://doi.org/10.1016/j.ydbio.2008.10.029

Legrand, J. M. D., & Hobbs, R. M. (2018). RNA processing in the male germline: Mechanisms and implications for fertility. Seminars in Cell and Developmental Biology, 79, 80–91. https://doi.org/10.1016/j.semcdb.2017.10.006

Monk, A. C., Siddall, N. A., Fraser, B., McLaughlin, E. A., & Hime, G. R. (2011). Differential roles of HOW in male and female Drosophila germline differentiation. PLoS ONE, 6(12), e28508–e28508. https://doi.org/10.1371/journal.pone.0028508

Monk, A. C., Siddall, N. A., Volk, T., Fraser, B., Quinn, L. M., McLaughlin, E. A., & Hime, G. R. (2010). HOW Is Required for Stem Cell Maintenance in the Drosophila Testis and for the Onset of Transit-Amplifying Divisions. Cell Stem Cell, 6(4), 348–360. https://doi.org/10.1016/j.stem.2010.02.016

Nabel-Rosen, H., Dorevitch, N., Reuveny, A., & Volk, T. (1999). The balance between two isoforms of the Drosophila RNA-binding protein How controls tendon cell differentiation. Molecular Cell, 4(4), 573–584. https://doi.org/10.1016/S1097-2765(00)80208-7

Nabel-Rosen, H., Volohonsky, G., Reuveny, A., Zaidel-Bar, R., & Volk, T. (2002). Two isoforms of the Drosophila RNA binding protein, How, act in opposing directions to regulate tendon cell differentiation. Developmental Cell, 2(2), 183–193. https://doi.org/10.1016/S1534-5807(01)00118-6

Nir, R., Grossman, R., Paroush, Z. e., & Volk, T. (2012). Phosphorylation of the <i>Drosophila melanogaster</i> RNA-binding protein HOW by MAPK/ERK enhances its dimerization and activity. PLoS Genetics, 8(3), e1002632–e1002632. https://doi.org/10.1371/journal.pgen.1002632

Poon, C. L. C., Zhang, X., Lin, J. I., Manning, S. A., & Harvey, K. F. (2012). Homeodomain-interacting protein kinase regulates Hippo pathway-dependent tissue growth. Current Biology, 22(17), 1587–1594. https://doi.org/10.1016/j.cub.2012.06.075

Ray, D., Kazan, H., Cook, K. B., Weirauch, M. T., Najafabadi, H. S., Li, X., … Hughes, T. R. (2013). A compendium of RNA-binding motifs for decoding gene regulation. Nature, 499(7457), 172–177. https://doi.org/10.1038/nature12311

Rodrigues, F., Thuma, L., & Klämbt, C. (2012). The regulation of glial-specific splicing of Neurexin IV requires HOW and Cdk12 activity. Development, 139(10), 1765–1776. https://doi.org/8223268

Teplova, M., Hafner, M., Teplov, D., Essig, K., Tuschl, T., & Patel, D. J. (2013). Structure-function studies of STAR family Quaking proteins bound to their in vivo RNA target sites. Genes & Development, 27(8), 928–940. https://doi.org/10.1101/gad.216531.113

Tettweiler, G., Blaquiere, J. A., Wray, N. B., & Verheyen, E. M. (2019). Hipk is required for JAK/STAT activity during development and tumorigenesis. PLoS One, 14(12), e0226856. https://doi.org/10.1371/journal.pone.0226856

Vernet, C., & Artzt, K. (1997). STAR, a gene family involved in signal transduction and activation of RNA. Trends in Genetics, 13(12), 479–484. https://doi.org/10.1016/S0168-9525(97)01269-9

