## Supplemental Figures for "Held out wings RNA binding activity in the cytoplasm during early spermatogenesis"

### Sup 1

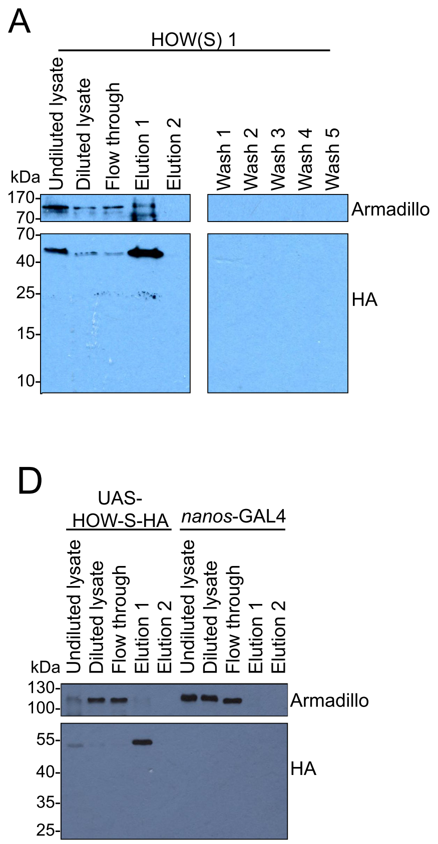


**Sup 1: Expression and pull-down of HA-tagged HOW(S) from cytoplasm of germ cells**
Western blots of HOW(S)-HA pull-down from both experimental *nanos*-GAL4>HOW(S)-HA flies (A-C) and parental control lines (D).

| Samples | | Raw reads | Pre-processing | | | Genome alignment | | | Transcriptome quasi-mapping | |
|---|---|---|---|---|---|---|---|---|---|---|
| | | | Poor quality | rRNA | tRNA | Unaligned | Aligned, unassigned | Aligned, assigned | Unmapped | Mapped |
| *nanos-* GAL4 | Total RNA | 53743743 | 11.46 | 39.27 | 0.0003 | 11.11 | 3.57 | 34.59 | 12.17 | 37.10 |
| | Pull down | 46955434 | 11.26 | 9.93 | 0.0018 | 25.17 | 6.82 | 46.83 | 29.43 | 49.39 |
| HOW(S) 1 | Total RNA | 53660667 | 10.47 | 66.03 | 0.0003 | 6.19 | 1.59 | 15.72 | 6.48 | 17.02 |
| | Pull down | 40300616 | 12.59 | 15.68 | 0.0003 | 13.02 | 8.08 | 50.63 | 20.89 | 50.85 |
| HOW(S) 2 | Total RNA | 55018237 | 11.80 | 42.54 | 0.0003 | 6.62 | 3.65 | 35.39 | 7.90 | 37.75 |
| | Pull down | 43895836 | 12.28 | 13.73 | 0.0018 | 12.42 | 8.39 | 53.18 | 19.27 | 54.72 |
| HOW(S) 3 | Total RNA | 52141872 | 11.29 | 42.25 | 0.0003 | 6.73 | 3.67 | 36.06 | 8.10 | 38.36 |
| | Pull down | 48005064 | 11.57 | 29.34 | 0.0014 | 10.75 | 5.76 | 42.58 | 13.96 | 45.12 |

**Sup Table 1: Read counts through RIP-seq pipeline by percentage.** Percentage of reads assigned in each step of the processing from the starting number of reads. ‘Unaligned’ refers to reads that were not aligned by Subread, ‘aligned, unassigned’ refers to reads that were aligned to the genome but not assigned to the ‘gene’ feature by featureCounts, and ‘aligned, assigned’ refers to reads that were both aligned to the genome and assigned the ‘gene’ feature by featureCounts.

### Sup 2

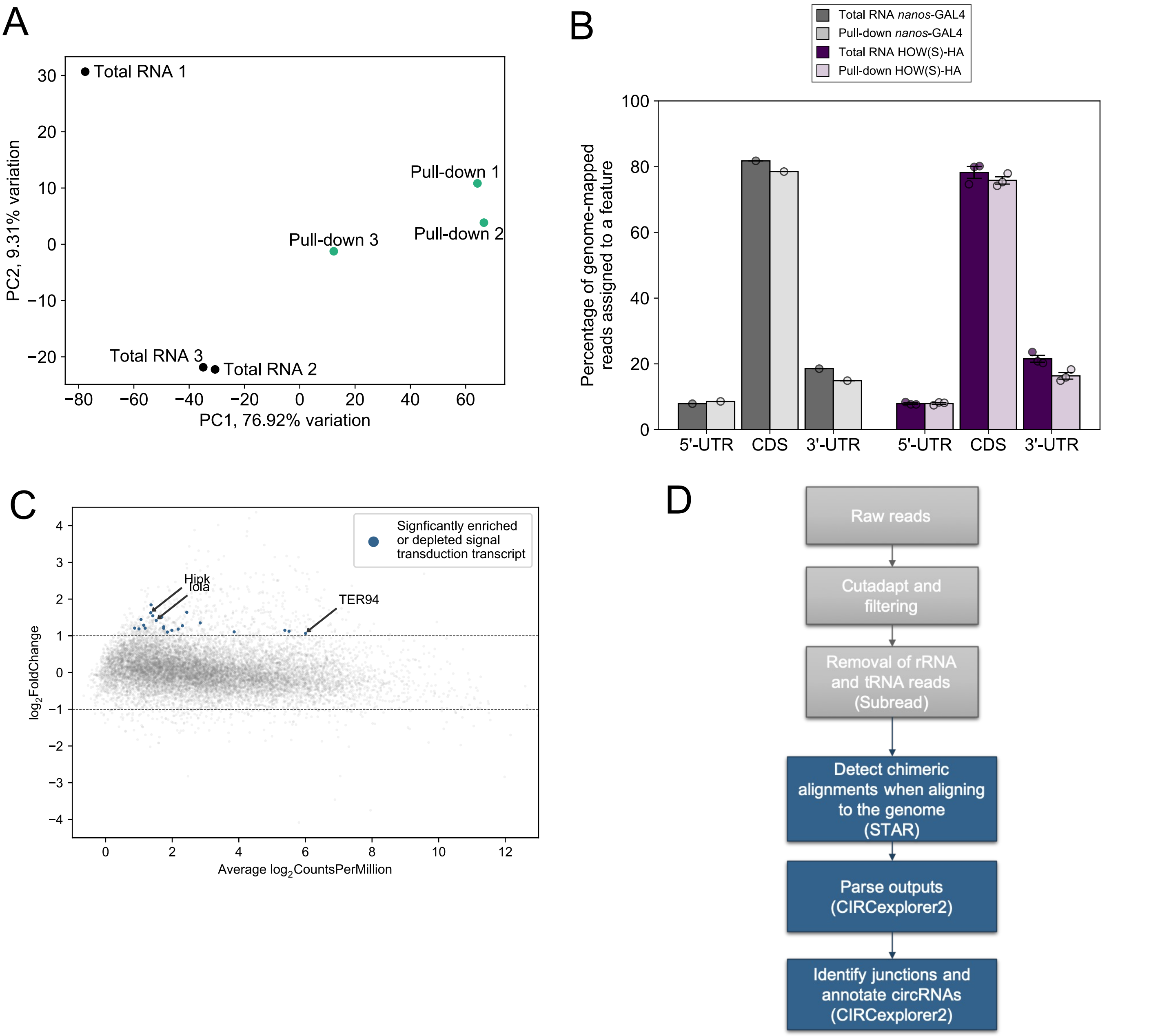


**Sup 2: Enrichment of HOW(S) bound RNA, with role in signal transduction**
A) Biplot from PCA using the $\log_2$(Counts Per Million) of the transcriptome quasi-mapped reads. B) Reads from HOW(S) and parental *nanos*-GAL4 RIP-Seq map across mRNA transcripts (5'-UTRs, CDSs and 3'-UTRs) at similar levels in the input and pull-down samples (error bars are SEM). C) Enriched mRNAs identified at transcript level whose protein products function in signal transduction marked in blue and 3 specific mRNAs of interest labelled. D) Pipeline for analysis of circRNAs bound by HOW(S).

| Location | Transcript ID | Name | nanos-GAL4 | | HOW(S)-HA 1 | | HOW(S)-HA 2 | | HOW(S)-HA 3 | |
|---|---|---|---|---|---|---|---|---|---|---|
| | | | Total | Pull-down | Total | Pull-down | Total | Pull-down | Total | Pull-down |
| X:12411364-12411589 | FBtr0347270 | CR32652 | 63 | 129 | 74 | 82 | 102 | 69 | 89 | 31 |
| 2R:17275409-17276063 | FBtr0310346 | muscleblind | 57 | 45 | 26 | 53 | 30 | 45 | 19 | 9 |
| 2R:21769766-21769904 | FBtr0300226 | CG30395 | 65 | 44 | 24 | 59 | 33 | 12 | 8 | 73 |
| 2R:24771724-24772795 | FBtr0072433 | Eps-15 | 3 | 9 | 12 | 4 | 7 | 1 | 6 | 2 |
| 4:1022241-1023680 | FBtr0309865 | Plexin A | 3 | 9 | 3 | 5 | 9 | 10 | 12 | 2 |
| 2L:20648188-20648248 | FBtr0300249 | Uhg3 | 1 | 5 | 4 | 5 | 9 | 1 | 8 | 7 |

**Sup Table 2: circRNAs from HOW(S)-HA RIP-Seq**
The 6 circRNAs with ≥ 9 reads aligned in at least 1 of the 8 samples. Those highlighted in blue contain the (A/G/U)CUAAC motif within them.

### Sup 3

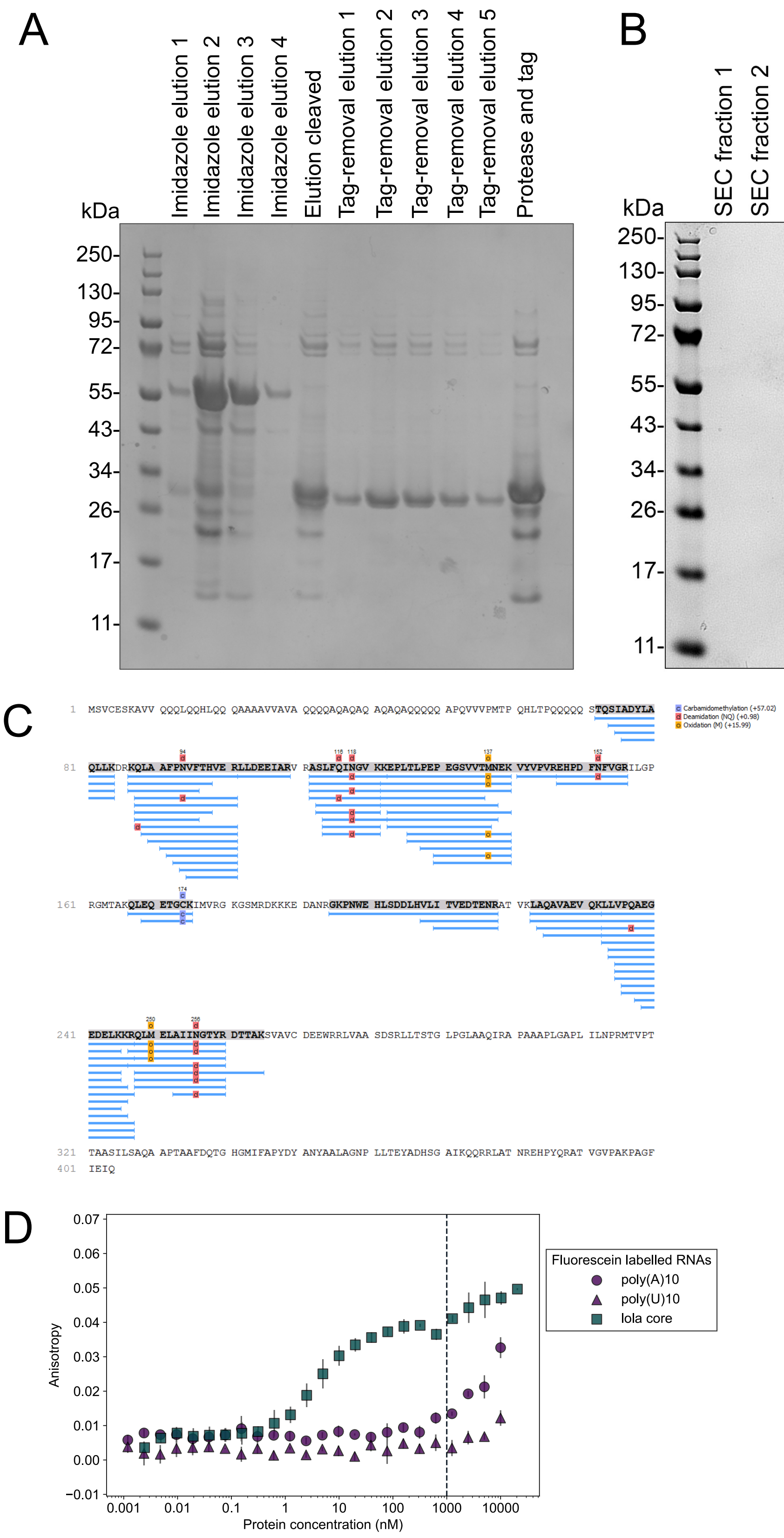


**Sup 3: HOW KH domain binds to HOW(S) targets with high affinity and specificity**
A-B) Coomassie stained gels from the purification of HOW-STAR domain. A) $Ni^{2+}$ affinity chromatography to purify $His_6$-GST tagged HOW-STAR. Imidazole elutions were combined and cleaved overnight (‘elution cleaved’) and the tag and protease were removed by another round of $Ni^{2+}$ affinity chromatography. B) Further purification of HOW-STAR performed by size exclusion chromatography. C) Mass spectrometry of purified HOW-STAR shows coverage across residues 72-265 (STAR domain). D) FA of HOW-STAR with 3 RNA oligos. Above 1000 nM of protein suspected non-specific binding occurs.
