## Supplemental Methods for "Held out wings RNA binding activity in the cytoplasm during early spermatogenesis"

**Immunofluorescence**

0-3 day old unmated male flies were anaesthetised and dissected in 1X PBS. The following fixing, washing and staining steps were carried out with samples rotating. Testes were fixed with 4% paraformaldehyde in PBS for 40 min at room temperature (RT). Washed with 1X PBX (Sup Table 3) three times, blocked with blocking buffer (Sup Table 3) for 1 hr at RT. Primary antibodies were used in 1X PBX and applied for 2 hr at RT or overnight at 4 °C. Testes were washed three times and then incubated with secondary antibodies for 2 hours at RT or overnight at 4 °C. This process was repeated for the next primary and secondary antibody pair. Samples were mounted with Vectashield antifade mounting medium with DAPI (Vector Laboratories). All antibodies used are described in Sup Table 4. Slides were imaged using a Zeiss LSM880 Upright Confocal Microscope with the 40X oil-immersion objective and Zen imaging software. The lasers used were Argon 488, DPSS 561 nm, and Diode 405 nm.

**Ribonucleoprotein immunoprecipitation**

The reproductive systems of 0-3 day old male flies (mated and unmated) were dissected in 1X PBS with 30 U/mL RNasin Plus RNase inhibitor (Promega). After 10 pairs of testes were dissected, tissue was snap frozen in liquid nitrogen. At least 1000 pairs of testes were collected per sample. Tissue was homogenised with an equal volume of RIP lysis buffer (Sup Table 3) with a micropestle and then with 23 gauge needle. Lysates were incubated on ice for 30 min, inverted halfway, and then snap frozen. Samples were thawed and centrifuged at 15,000 xg for 15 minutes at 4°C. 20 µL or 10% of supernatants (whichever was less) was set aside and snap frozen as the ‘input’ samples.

180 µL of anti-HA beads (Pierce, #88837) per sample were used and washed 5 times in ice-cold NT2 buffer (Sup Table 3) prior to usage. Beads were resuspended in NT2 buffer at 5.5 times the volume of sample and 0.2 U/μL of RNase inhibitor and EDTA pH 8 to a final concentration of 20 mM were added. Beads and lysates were combined and incubated while tumbling for 1 hr at RT. After incubation the supernatant was taken as the ‘flow through’ samples and snap frozen. Beads were then washed five times in 1 mL NT2 buffer.

A quarter of the beads were eluted for protein samples, these were boiled at 95 °C for 10 min in 25 µL of 2X protein sample buffer (Sup Table 3). These samples were analysed by western blotting. The antibodies used for western blots are described in Sup Table 5.

The remaining three quarters of the beads went through RNA elution and extraction. First, the elution samples were treated with 20 mg/mL proteinase K at 65 °C for 35 min. Then supernatants from this step, alongside proteinase K treated ‘input’ samples, were treated with 3 times their volume in Trizol for 5 min at RT. 0.15% (v/v) chloroform was added and tubes were vigorously shaken for 15 sec, incubated for 3 min at RT before being centrifuged at 12,000 xg for 15 min at 4 °C. The aqueous phase was collected and 1 volume of isopropanol was added along with 1 µL glycoblue and 0.3 M NaCl. The RNA was precipitated at -80 ºC for at least 3 hours. After precipitation, samples were centrifuged at 13,300 xg for 20 min at 4 ºC. The pellet was washed twice with 70% ethanol with 5 min centrifuge spin in between. After the final spin the supernatant was removed and the pellet was left to air-dry for 10–15 minutes. The pellet was resuspended in 18 µL nuclease-free water.

**Sup Table 3 – Buffers and solutions used.** All descriptions are for 1X solutions unless stated otherwise.

| **Buffer name** | **Buffer components** |
| --- | --- |
| PBX | 0.1% (v/v) Triton X-100, 0.5% (v/v) normal goat serum (Thermo Fisher), in 1X phosphate buffered saline (PBS) |
| Blocking buffer | 0.1% Triton X-100, 2% (v/v) normal goat serum, in 1X PBS |
| NT2 buffer | 50 mM Tris-HCl pH 7.4, 150 mM NaCl, 1 mM MgCl2, 0.05% (v/v) IGEPAL |
| PBS-T | 0.1% Tween 20, in 1X PBS |
| 2X protein sample buffer | 125 mM Tris-HCl pH 6.8, 5% (w/v) SDS, 25% (v/v) glycerol, 10% (v/v) β-mercaptoethanol, 0.004% (w/v) bromophenol blue |
| RIP lysis buffer | 50 mM Tris-HCl pH 8, 150 mM NaCl, 10 mM MgCl2, cOmplete mini protease inhibitor cocktail (Roche), 1% (v/v) IGEPAL, 24 U/mL Turbo DNase (Thermo Fisher), 30 U/mL RNasin Plus RNase inhibitor (Promega) |
| SEC buffer | 25 mM HEPES pH 7.6, 150 mM NaCl and 1 mM TCEP |
| RNA binding buffer | 20 mM Tris-HCl pH 7.5, 150 mM NaCl, 0.01% Triton X-100 |

**Sup Table 4 – Antibodies used for immunofluorescence.** Primary antibodies are listed in the top half of the table and secondary antibodies in the bottom half.

| **Antibody** | **Additional information** | **Stock concentration (µg/mL)** | **Dilution used** | **Supplier**  **(catalogue reference)** |
| --- | --- | --- | --- | --- |
| Vasa | Rat IgM  Monoclonal  Hybridoma supernatant | 44 | 1:200 | DSHB  (anti-vasa) |
| HA | Mouse IgG2b  Monoclonal  Ascites fluid | 400 | 1:100 | Roche  (12CA5) |
| Goat anti-rat IgM | Alexa Fluor 488  Polyclonal  Affinity purified | 2000 | 1:400 | Thermo Fisher  (A-21212) |
| Goat anti-mouse IgG2b | Alexa Fluor 594  Polyclonal  Affinity purified | 2000 | 1:400 | Thermo Fisher  (A-21145) |

**Sup Table 5 – Antibodies used for western blotting.** Primary antibodies are listed in the top half of the table and secondary antibodies in the bottom half.

| **Antibody** | **Additional information** | **Stock concentration (µg/mL)** | **Dilution used** | **Supplier**  **(catalogue reference)** |
| --- | --- | --- | --- | --- |
| Armadillo | Mouse IgG2a  Monoclonal  Hybridoma supernatant | 27 | 1:1000 | DSHB  (N2 7A1) |
| HA | Rabbit IgG  Polyclonal  Affinity purified | 1000 | 1:5000 | Abcam  (ab9110) |
| Horse anti-mouse IgG | HRP-linked | 153 | 1:5000 | Cell Signalling Technology  (7076S) |
| Goat anti-rabbit IgG | HRP-linked | 65.7 | 1:5000 | Cell Signalling Technology  (7074S) |

**Sequencing and computational analysis**

*Sequencing*

The RNA from the lysates and elutions of three HOW(S)-HA samples and the *nanos*-GAL4 parental control was prepared using the Ribo-Zero rRNA Removal Kit (Illumina) followed by the TruSeq Stranded Total RNA Library Prep (Illumina). 100 ng of each sample was pooled and sequenced on the same lane on the NextSeq 500 Illumina sequencer using the High Output Kit v2.5 (75 Cycles), i.e. 75 bp single-end sequencing.

*Filtering*

The adapter sequence (AGATCGGAAGAGCACACGTCTGAACTCCAGTCAC) was trimmed from reads with Cutadapt (version 1.1) (Martin, 2011). Reads were filtered out if the quality score was below than 20 for 10% or more of the read. Filtering was done using the ‘Filter by quality’ tool in Galaxy (version 1.0.2) (Gordon, 2010). Next, rRNA and tRNA reads were removed using Subread (version 2.0.0) (Liao et al., 2013). The tRNA fasta file was from release 6.22 of the *D. melanogaster* genome. The rRNA fasta file was from RiboGalaxy’s Shared Data library (UUID: 0f0983aa-3afb-4b5f-a417-23593a3df1ef).

*Transcriptome mapping*

Transcriptome indexing and quasi-mapping was carried out using Salmon (version 0.14.2) (Patro et al., 2017). A decoy-aware transcriptome was indexed with a pre-computed decoy sequence file provided by the Salmon developers. The transcriptome and decoy files used were based on the *D. melanogaster* genome from Ensembl release 97, which corresponds to FlyBase’s release 6.22. The k-mer size selected for indexing the transcriptome was 31. The reads remaining after filtering were quasi-mapped to the transcriptome with library parameter (-l) set to SR to correspond with the reverse stranded library type. The k-mer parameter (-k) was set to 31.

*Differential transcript enrichment*

The following analysis was executed in RStudio, version 0.99.486, with R version 3.6.2 (Team, 2020). Transcript counts from Salmon were imported using the tximport package (version 1.14.0) (Soneson et al., 2015) using txOut=TRUE and countsFromAbundance="scaledTPM" while importing. Differential enrichment was carried out with the edgeR package (version 3.28.0) (McCarthy et al., 2012; Robinson et al., 2010). Low expression transcripts were filtered using the filterByExprfunction, libraries were normalised using calcNormFactors.

Two factors were defined for the HOW(S)-HA samples: condition and pairing. Condition referred to whether a sample was a total RNA or a pull-down RNA sample. Pairing referred to the three pairs of total and pull-down RNA samples. Thus, the design matrix was submitted as follows: model.matrix(~sample$condition

+ sample$pair). One factor was defined for the *nanos*-GAL4 samples to differenti-

ate between the total RNA and pull-down RNA. The design matrix was submitted as:

model.matrix(~group). For the HOW(S)-HA samples, dispersion was estimated with the design matrix taken into account. Then testing for differential genes and transcripts was carried out with the quasi-likelihood F-test, as per the edgeR manual instructions. For the *nanos*-GAL4 samples, the exact test was used with the dispersion set as the square of the biological coefficient of variation (BCV). The BCV was set as the square root of the common dispersion from the HOW(S) data.

Finally, transcripts from the HOW(S)-HA RIP-seq data were classed as significantly enriched or depleted if they had an adjusted *p*-value < 0.05, had a log_2_(Fold Change) above 1 or below -1, and if it did not meet these same thresholds in the *nanos*-GAL4 data.

*Gene ontology*

Gene ontology analysis was carried out in Gene Ontology enRIchment anaLysis and visuaLizAtion tool (GOrilla) (Eden et al., 2007; Eden et al., 2009). The running mode used two unranked lists of genes: 1) enriched transcript list converted to their gene IDs, 2) transcripts that were not filtered out by edgeR’s filterByExpr function converted to their gene IDs. *D. melanogaster* genes under the GO term ‘signal transduction’ (GO:0007165) were accessed via FlyBase’s controlled vocabulary tool.

*Motif enrichment*

Discriminative Regular Expression Motif Elicitation (DREME) from MEME Suite (MEME version 5.1.0, with Python version 2.7.15) was used to carry out the motif enrichment analysis on 5’ and 3’-UTRs (Bailey, 2011). Control sequences were generated from the list of transcripts that were not filtered out by edgeR’s filterByExpr function. DREME was implemented using the -norc flag so only the strand given was searched and not the complimentary sequences, -rna flag was used to indicate the sequences were of RNA not DNA, and -m was set to 25 to stop searching after 25 motifs had been found.

*Principal Component Analysis*

Principal component analysis (PCA) was performed using the PCAtools (version 1.2.0) (Blighe, 2019) package in R. PCA was carried out on the triplicate HOW(S)- HA pull-down samples with the log_2_(Counts Per Million) from the transcript-level data, as calculated by the edgeR package in section 2.6.3.4. The pca function was used with the removeVar parameter set to 0.1, this removes the lower 10% of variables based on variance. The biplot function was used to generate a graph of PC1 against PC2.

*Read summarisation*

To summarise the genomic features, the filtered reads were aligned to the *D. melanogaster* genome release 6.22 using Subread. The type parameter (-t) was set to 0 to indicate RNA-seq reads and the remaining parameters were left at their default settings. featureCounts was used to count reads to the following features: ‘gene’, ‘5UTR’, ‘CDS’ and ‘3UTR’ (Liao et al., 2014). The GTF file used for this was from D. melanogaster genome release 6.22, parameter -s was set to 1 to indicated the library preparation results in a stranded library, -g was used to group features into the gene ID meta-feature. The remaining parameters were left at their default settings. Count tables from the ‘gene’ feature were used in Sup Table 1.

*Circular RNA alignment and annotation*

An annotation file was retrieved from Ensembl’s release 97 for *D. melanogaster* in the gene transfer format (GTF) (Zerbino et al., 2018), which corresponds to FlyBase’s release 6.22. All rows with ‘gene’ in the third column were removed with a custom script, this is the modified GTF file. The file was then converted to the GenePred table format using the gtfToGenePred function from the ucsc-gtftogenepred package (version 366). Finally, a new first column was added with ‘ens97’ added to every row with a custom script, this is the modified GenePred file.

Spliced Transcripts Alignment to a Reference (STAR; version 2.7.3a) was used to align the filtered RIP-seq reads to release 6.22 of the *D. melanogaster* genome (Dobin et al., 2013). First, the genome was indexed using by setting the --runmode parameter to genomeGenerate. The --sjdbOverhang parameter was set to 75, --genomeSAindexNbases was set to 13, the modified GTF file described above was used as the input annotation file with FlyBase’s all chromosome fasta file from release 6.22 of the *D. melanogaster* genome. All other parameters were left at their default settings. Reads were aligned to the genome by setting the --runmode parameter to alignReads. The --chimSegmentMin parameter was set to 15 and --quantMode set to GeneCounts, all other parameters left at their default settings.

CIRCexplorer2 (version 2.3.8) was used to identify circRNAs from the STAR aligned files (Zhang et al., 2016). First, the Chimeric.out.junction output files from STAR were parsed using CIRCexplorer2’s parse function, with the -t parameter set to STAR. The back spliced junction.bed output files were then annotated using STAR’s annotate function with the modified GenePred file and FlyBase’s all chromo- some fasta file from release 6.22 of the *D. melanogaster* genome. The values in the circularRNA_known.txt output files were used as the final counts for all circRNAs identified.

**Protein expression and purification**

Residues 72–266 of *D. melanogaster* HOW (HOW-PB; FBpp0083576), the STAR domain, was codon optimised and ordered from Genewiz and cloned into the pOPIN-J vector (Berrow et al., 2007). The His_6_-GST-STAR domain was expressed in BL21(DE3) cells overnight at 18 ºC. The fusion protein was affinity purified with a HisTrap column (GE Healthcare), followed by cleavage with 3C protease overnight. The cleavage products were applied to a HisTrap column to remove the His_6_-GST tag. The cleaved STAR domain was further purified by size exclusion chromatography (SEC) using a 26/600 Superdex 75 column and SEC buffer (Sup Table 3).

**Fluorescence anisotropy**

RNA oligonucleotides were synthesised by Integrated DNA Technologies. Oligos were 3’ labelled with 6-carboxyfluorescein and all oligos were desalted after synthesis. Binding assays between fluorescein labelled oligos and purified STAR protein were carried out in triplicate in black 384-well OptiPlates (Perkin Elmer), control wells (with no RNA) were carried out once. 20 μL of RNA binding buffer (Sup Table 3) was added to every well. 20 μL of 4 nM protein was added to the first column and titrated across each row. Finally, 20 μL of 5 nM fluorescein labelled RNA oligo was added to the appropriate rows, and for the control rows 20 μL of RNA binding buffer was added instead. The plate was left to equilibrate for at least 45 minutes. Then data was collected on a Spark 10M Multimode Microplate Reader (Tecan) with a 485 nm (20 nm bandwidth) excitation filter, and parallel (S) and perpendicular (P) channel emission filters at 535 nm (25 nm bandwidth).

Anisotropy values were calculated using the emission values for S and P signals (corrected with the control values from protein only wells) with the following equation:

$$Anistropy= \frac{S-P}{S+2P}$$

The logistic function in OriginPro 2020 V2 was first used to determine the theoretical minimum and maximum anisotropy values (A_1_ and A_2_, respectively), using the equation below:

$$y=A_{2}+ \frac{A_{1}-A_{2}}{1+\left( \frac{x}{x_{0}} \right)^{p}}$$

Where $y$ is either anisotropy or fraction bound, $x$ is protein concentration, $x_{0}$ is the dissociation constant and $p$ is the Hill coefficient. A_1_ and A_2_ were used to calculate the fraction of RNA bound. The logistic function was used again to fit a curve for the fraction bound data. The dissociation constant (x_0_) from these curves was reported as the apparent K_D_.

**References**

Bailey, T. L. (2011). DREME: motif discovery in transcription factor ChIP-seq data. *Bioinformatics*, *27*(12), 1653-1659. <https://doi.org/10.1093/bioinformatics/btr261>

Berrow, N. S., Alderton, D., Sainsbury, S., Nettleship, J., Assenberg, R., Rahman, N., . . . Owens, R. J. (2007). A versatile ligation-independent cloning method suitable for high-throughput expression screening applications. *Nucleic Acids Res*, *35*(6), e45. <https://doi.org/10.1093/nar/gkm047>

Blighe, K. (2019). PCAtools: PCAtools: Everything Principal Components Analysis. In.

Dobin, A., Davis, C. A., Schlesinger, F., Drenkow, J., Zaleski, C., Jha, S., . . . Gingeras, T. R. (2013). STAR: ultrafast universal RNA-seq aligner. *Bioinformatics*, *29*(1), 15-21. <https://doi.org/10.1093/bioinformatics/bts635>

Eden, E., Lipson, D., Yogev, S., & Yakhini, Z. (2007). Discovering Motifs in Ranked Lists of DNA Sequences. *PLoS Computational Biology*, *3*(3), e39-e39. <https://doi.org/10.1371/journal.pcbi.0030039>

Eden, E., Navon, R., Steinfeld, I., Lipson, D., & Yakhini, Z. (2009). GOrilla: A tool for discovery and visualization of enriched GO terms in ranked gene lists. *BMC Bioinformatics*, *10*(1), 48-48. <https://doi.org/10.1186/1471-2105-10-48>

Gordon, A. (2010). FASTQ/A short-reads pre-processing tools. In.

Liao, Y., Smyth, G. K., & Shi, W. (2013). The Subread aligner: Fast, accurate and scalable read mapping by seed-and-vote. *Nucleic Acids Research*, *41*(10). <https://doi.org/10.1093/nar/gkt214>

Liao, Y., Smyth, G. K., & Shi, W. (2014). featureCounts: an efficient general purpose program for assigning sequence reads to genomic features. *Bioinformatics*, *30*(7), 923-930. <https://doi.org/10.1093/bioinformatics/btt656>

Martin, M. (2011). Cutadapt removes adapter sequences from high-throughput sequencing reads. *EMBnet.journal*, *17*(1), 10-10. <https://doi.org/10.14806/ej.17.1.200>

McCarthy, D. J., Chen, Y., & Smyth, G. K. (2012). Differential expression analysis of multifactor RNA-Seq experiments with respect to biological variation. *Nucleic Acids Research*, *40*(10), 4288-4297. <https://doi.org/10.1093/nar/gks042>

Patro, R., Duggal, G., Love, M. I., Irizarry, R. A., & Kingsford, C. (2017). Salmon provides fast and bias-aware quantification of transcript expression. *Nature Methods*, *14*(4), 417-419. <https://doi.org/10.1038/nmeth.4197>

Robinson, M. D., McCarthy, D. J., & Smyth, G. K. (2010). edgeR: a Bioconductor package for differential expression analysis of digital gene expression data. *Bioinformatics*, *26*(1), 139-140. <https://doi.org/10.1093/bioinformatics/btp616>

Soneson, C., Love, M. I., & Robinson, M. D. (2015). Differential analyses for RNA-seq: transcript-level estimates improve gene-level inferences. *F1000Research*, *4*, 1521-1521. <https://doi.org/10.12688/f1000research.7563.1>

Team, R. S. (2020). RStudio: Integrated Development Environment for R. In. Boston, MA.

Zerbino, D. R., Achuthan, P., Akanni, W., Amode, M. R., Barrell, D., Bhai, J., . . . Flicek, P. (2018). Ensembl 2018. *Nucleic Acids Research*, *46*(D1), D754-D761. <https://doi.org/10.1093/nar/gkx1098>

Zhang, X. O., Dong, R., Zhang, Y., Zhang, J. L., Luo, Z., Zhang, J., . . . Yang, L. (2016). Diverse alternative back-splicing and alternative splicing landscape of circular RNAs. *Genome Research*, *26*(9), 1277-1287. <https://doi.org/10.1101/gr.202895.115>
